# Resting-state functional connectivity after creativity training with music composing

**DOI:** 10.64898/2026.01.29.701494

**Authors:** Anna Arkhipova, Pavel Hok, Markéta Trnečková, Gabriela Žatková, Vít Zouhar, Petr Hluštík

## Abstract

Creativity is one of the unique cognitive constructs in human beings and its neurobiological correlates are one of the current hot topics in neuroscience. The “Different Hearing” program (DHP) is an educational activity aimed at stimulating musical creativity by means of group composing in the classroom, alternative to the mainstream model of music education in Czechia. In our previous study, the data from task-related functional MRI with passive listening was analyzed. The results suggested that DHP training modified the response to diverse sound samples, differentially changing the engagement of functional networks known to be related to creative thinking, namely, increasing default mode network activation and decreasing activation of executive and salience networks. In the present study, we hypothesized that the DHP short-term (2 days) intense workshop would also induce changes in the resting-state networks that were significantly modified during task. To investigate it, seed-based, ROI-to-ROI resting-state functional connectivity and degree centrality analysis were performed on the acquired resting-state fMRI data. The results showed no significant group-by-time interaction.

## INTRODUCTION

Creativity is one of the most important abilities of human beings and methods and mechanisms to enhance creativity have fascinated researchers across a wide range of human activities (Arkhipova et al., 2026). Creativity has been defined as an ability to produce novel and useful ideas/works both in the public and among scholars (Runco and Jaeger, 2012). The Different Hearing program (DHP) is an educational initiative designed to stimulate musical creativity through collaborative composition in classroom settings, offering an alternative to the dominant model of music education in Czechia (Zouhar and Medek, 2010). In our previous study (Arkhipova et al., 2021), the data from task-related fMRI with passive listening was analyzed. The results suggested that short term (2days) DHP training modified the response to diverse sound samples, differentially changing the engagement of functional networks known to be related to creative thinking, namely, increasing default mode network (DMN) activation and decreasing activation of executive networks (EN) and salience network (SN).

An association between DMN and high creativity trait was suggested by several fMRI studies (Takeuchi et al., 2012; Beaty et al., 2014; Chen et al., 2014; Li et al., 2016; Zhu et al., 2017): positive correlation with resting-state functional connectivity (rsFC) within DMN (Takeuchi et al., 2012; Chen et al., 2014) or between the EN and DMN regions (Beaty et al., 2014; Zhu et al., 2017); negative correlations with rsFC within DMN (Li et al., 2016; Zhu et al., 2017).

Specifically for musical creativity, a cross-sectional study by Bashwiner et al. (2020) showed significant correlation with rsFC including the regions of DMN, EN and SN. On the other hand, previous interventional studies using rsfMRI in verbal creativity training also point to the central role of the DMN (Wei et al., 2014; Fink et al., 2018; Sun et al., 2020) and its interactions with middle temporal gyrus (Wei et al., 2014) and dorsal anterior cingulate cortex (ACC, a central hub of the SN - Sun et al., 2020). Also, a change in rsFC within the EN was observed by Fink et al. (2018). The DMN has therefore been proposed to underlie high creative thinking by hosting spontaneous cognitive processes that interact with high executive functions represented by the EN (Arkhipova et al., 2026). Recently, graph-theoretical analyses have been also used to identify networks in rsfMRI data associated with creative expertise, showing for example, higher local efficiency within the DMN (Chen et al., 2019), or lower centrality in the primary visual cortex was in experts (Orwig et al., 2023). However, we are not aware of any studies which investigated changes in rsFC or degree centrality after short-term musical creativity training.

In this study, we have investigated the rsFC changes after 2-day music creativity training with the hypothesis: 1) DHP workshop would induce FC pattern changes in the DMN, EN and SN; 2) change in degree centrality in the corresponding networks would be observed after the workshop.

## METHODS

### Subjects

Forty-six healthy students from the Faculty of Education at Palacký University Olomouc (40 females and 6 males, mean age 21.6 ± 1.4) were randomly assigned into two groups. Both groups were matched in terms of music education, according to computed results from self-reported questionnaires. Four participants were left-handed, two were ambidextrous, and 41 were right-handed as assessed by the Edinburgh handedness inventory. The study was carried out in accordance with World Medical Association Declaration of Helsinki. Written informed consent was obtained from all participants prior to their inclusion in the study and the study was approved by the Ethics Committee of the Faculty of Education at Palacký University in Olomouc, approval number 02/2017.

### Study protocol

All participants underwent 2 functional MRI examinations 8 days apart on a 3T scanner (Prisma, Siemens Healthcare, Erlangen, Germany) including structural, resting-state fMRI (rs-fMRI) and task-related data. Between the fMRI examinations, the active group (22 participants) participated in the DH workshop over two days and the control group (24 participants) continued normal daily activities and student life without any special training. Further details of protocol can be cited from the previous study (Arkhipova et al., 2021).

### Intervention: the “Different Hearing” Program Workshop

The aim of DHP is to stimulate and enhance participants’ musical creativity through group learning to compose music in the classroom. The program started in 2001 at the Department of Music Education, Palacký University Olomouc, Czechia, as an alternative to the common model of music education. For our study, the DHP workshop lasted 10.5 h, split into two consecutive days. During the workshop, the participants learned to use common objects and their bodies to make sounds, also employed natural and environmental sounds as the “building blocks” in composing music. The team-wise creative process ended by writing down a graphical score and finally performing the composition in front of an audience. For detailed description of the workshop, see Arkhipova et al., 2021.

### Behavioral task

Behavioral data were collected during two runs of task-based fMRI acquisition during listening to musical and non-musical sound samples from five different classes, i.e., Classic music, Modern music, DHP samples (composed and performed by previous participants during the workshops in the past), Nature sounds, and Industrial noises (for the detailed sound selection, see Arkhipova et al., 2021). During each imaging run, fifteen different samples of 15s long were played through MR-compatible headphones in a counterbalanced order across subjects. Participants were asked to keep their eyes open to watch a fixation cross during the listening phase, instructed to press one of two buttons (like/not like) after listening to each sample as soon as the question “Did you like the sample?” (in Czech) appeared on screen (displayed for 4 s following the sound sample offset).

### Resting-state fMRI data acquisition

MRI examinations were performed twice with an 8-day interval for all subjects, using a 3T scanner (Siemens Prisma, Erlangen, Germany) with a standard 64-channel head and neck coil in the Multimodal and Functional Imaging Laboratory (MAFIL), Central European Institute of Technology (CEITEC) in Brno. The subject’s head was immobilized with cushions to assure maximum comfort and minimize head motion. The MRI protocol included resting-state BOLD fMRI data acquisition (T_2_∗ -weighted echo-planar imaging; 48 slices, 3-mm slice thickness; repetition time/echo time = 780/35 ms; flip angle 45∘; field of view = 192 mm; matrix 64 × 64; 465 volumes; time of acquisition = 6:00 minutes), subjects had their eyes closed. Gradient-echo phase and magnitude fieldmap images were acquired to allow correction of the echo-planar imaging distortions. A high resolution T1-weighted structural image was acquired using Magnetization-Prepared rapid Gradient-Echo (MPRAGE) sequence for anatomical reference. Task-related fMRI data (reported in our previous study) were obtained after resting-state fMRI imaging data. Heart rate (pulse oximetry) and respiration (respiratory belt) were monitored during BOLD scanning.

### Analysis of Behavioral data

The effect of time/session on subjective like/not like response to individual stimulus classes was tested within subject using Wilcoxon signed rank tests in the active and control groups.

### MRI Data Pre-processing

The rs-fMRI data were processed and analyzed using CONN Functional Connectivity Toolbox v. 21a (Whitfield-Gabrieli and Nieto-Castanon, 2012) for SPM 12 (v 7771, https://www.fil.ion.ucl.ac.uk/spm/software/spm12/) running under Matlab v. R2017b (https://www.mathworks.com/products/matlab.html). The pre-processing pipeline included motion correction, correction of susceptibility-induced distortions, motion outlier detection (global signal [GS] z-value threshold =3 and subject-motion threshold = 0.5 mm), spatial smoothing with full width at half maximum (FWHM) = 8 mm, and normalization to standard space. Next, a built-in anatomical component-based denoising procedure (aCompCor) and band-pass filtering within the frequency range 0.008 Hz – 0.09 Hz were applied. Additionally, 4 participants were identified as outliers (defined as values greater than Q3 + 1.5 * interquartile range) for number of invalid scans (threshold 6.0% of all scans) and maximum volume-to-volume head displacement >2.11 mm (3 in active and 1 in control group). After the subject exclusion, the distribution of correlations between functional connectivity and data quality measures followed random normal distribution with the 95% match criterion (Morfini et al., 2023).

### Resting-state functional connectivity analysis

Bivariate rsFC was evaluated for 15 regions of interest (ROIs) based on network atlas implemented in CONN, including: 4 from DMN, 7 SAL ROIs and 4 FP = EN (Table 1). We used the networks that show significant activation changes in our previous task-related fMRI study as seed regions. First, a seed-based rsFC of each ROI per session was evaluated in each individual by extracting mean BOLD time series from the ROIs and calculating seed-to-voxel Pearson correlation coefficients for each voxel. Next, Fisher transform was applied, yielding z scores as a measure of rsFC. To test for a connectivity, change in the active group compared to control group, an analysis of variance (ANOVA) model using group and time as factors was used to search for group-by-time interaction with cluster-level threshold with cluster-forming p < 0.001 and false-discovery-rate error (FDR)-corrected p < 0.0033 (Bonferroni-corrected for the number of ROIs). Additionally, ROI-to-ROI rsFC was evaluated among 15 network ROIs, searching for group by time interaction with cluster-level threshold at FDR-corrected p < 0.05 (MPVA omnibus test) and p < 0.05 connection-level threshold.

**Table 1.** List of regions of interest. Table lists regions of interests (ROIs) used in the ROI analysis. Areas are extracted from network ROIs in CONN v. 21a (Whitfield-Gabrieli and Nieto-Castanon 2012).

| Number | Network | Region |
| --- | --- | --- |
| 1 | Default mode network | Medial Prefrontal Cortex |
| 2 |  | Lateral Parietal Cortex (Left) |
| 3 |  | Lateral Parietal Cortex (Right) |
| 4 |  | Posterior Cingulate Cortex (incl. Precuneus) |
| 5 | Salience network | Anterior Cingulate Cortex |
| 6 |  | Anterior Insula (Left) |
| 7 |  | Anterior Insula (Right) |
| 8 |  | Rostral Prefrontal Cortex (Left) |
| 9 |  | Rostral Prefrontal Cortex (Right) |
| 10 |  | Supramarginal Gyrus (Left) |
| 11 |  | Supramarginal Gyrus (Right) |
| 12 | Executive network (frontoparietal network) | Lateral Prefrontal Cortex (Left) |
| 13 |  | Lateral Prefrontal Cortex (Right) |
| 14 |  | Posterior Parietal Cortex (Left) |
| 15 |  | Posterior Parietal Cortex (Right) |

## Degree centrality analysis

To obtain a whole-brain degree-centrality map with reasonable dimensionality, subject-specific functional connectivity matrices containing Fisher-transformed Pearson’s r coefficients were computed in CONN for 6,975 6-mm large voxels created by resampling the common gray matter mask. Degree centrality was then calculated for each subject using a brain connectivity toolbox (BCT, https://sites.google.com/a/brain-connectivity-toolbox.net/bct/) with 10% link density. Regional degree centrality was extracted by averaging nodal degree from 15 ROIs representing DMN, salience and executive networks, implemented in CONN. Pre-post differences in degree centrality within those ROIs were then compared using two-sample t-tests with Bonferroni-correct p < 0.0041 considered significant. To search for unsuspected group effects on degree centrality, an exploratory whole-brain analysis was carried out in randomize, part of FSL, v. 6.0.5 (https://fsl.fmrib.ox.ac.uk/fsl/docs/) using two-sample t-test contrast to compare pre-post degree centrality changes, yielding family-wise error-corrected significance maps thresholded at p < 0.05 after threshold-free cluster enhancement with 10,000 permutations.

## RESULTS

### Behavioral Data

Comparison of responses to sound samples showed that favorable feelings toward DHP, Modern music and non-musical sound samples (Nature and Industrial) significantly increased only in the active group (p < 0.05, Wilcoxon signed rank tests). The change for the DHP sound samples was most robust and survived optional Bonferroni correction for multiple testing.

### Resting-state fMRI functional connectivity

The seed-based connectivity analysis of rs-fMRI data showed no significant seeds between groups across time (see Table 2 for uncorrected results). No significant ROI-to-ROI connectivity pattern change was observed (see Figure 1 and Table 3 for uncorrected results).

**Table 2.** Uncorrected results of seed-based analysis. Table showing clusters with uncorrected cluster-size threshold of p < 0.05, that did not pass the FDR threshold (p < 0.0033). Cluster size is provided in voxels in the 2-mm standard space. Abbreviations: AI, Anterior Insula; FDR, false discovery rate; L, left; LP, Lateral Parietal Cortex; R, right; rPFC, Rostral Prefrontal Cortex; SMG, Supramarginal Gyrus; unc, uncorrected.

| Seed | Cluster (x, y, z) | Size | Size p-FDR | Size p-unc | Peak p-unc |
| --- | --- | --- | --- | --- | --- |
| LP (L) | +38 +20 +40 | 83 | 0.335511 | 0.023965 | 0.000024 |
| AI (R) | +46 -02 -10 | 83 | 0.422855 | 0.022256 | 0.000001 |
|  | +56 +28 +26 | 59 | 0.458635 | 0.048277 | 0.000051 |
| rPFC (R) | +34 +14 +44 | 167 | 0.058758 | 0.002422 | 0.000009 |
|  | -30 +16 +42 | 158 | 0.058758 | 0.003013 | 0.000002 |
|  | +60 -42 -26 | 102 | 0.170173 | 0.013090 | 0.000003 |
|  | -44 -72 -52 | 87 | 0.197431 | 0.020249 | 0.000078 |
|  | -20 -34 -20 | 64 | 0.299413 | 0.041676 | 0.000012 |
|  | +46 -60 +32 | 61 | 0.299413 | 0.046064 | 0.000219 |
| SMG (L) | -22 -62 -50 | 71 | 0.446394 | 0.032688 | 0.000113 |
| SMG (R) | -54 +28 +06 | 151 | 0.099577 | 0.003830 | 0.000022 |
|  | -08 -80 -22 | 86 | 0.213392 | 0.021849 | 0.000029 |
|  | -36 -36 +66 | 82 | 0.213392 | 0.024622 | 0.000135 |
|  | +36 -44 +60 | 62 | 0.300466 | 0.046225 | 0.000068 |

**Table 3.** Uncorrected results of the ROI-to-ROI analysis. Table showing the network connections among 15 ROIs at uncorrected connection-level p < 0.05, that did not pass the cluster-level threshold at FDR-corrected p < 0.05. Abbreviations: ACC, Anterior Cingulate Cortex; DMN, Default Mode Network; EN, Executive Network; FDR, false discovery rate; L, left; LP, Lateral Parietal Cortex; LPFC; Lateral Prefrontal Cortex, mPFC, Medial Prefrontal Cortex; PCC, Posterior Cingulate Cortex; PPC, Posterior Parietal Cortex; R, right; ROI, region of interest; rPFC, Rostral Prefrontal Cortex; SN, Salience Network.

| Connections | Statistic | p-uncorrected | p-FDR |
| --- | --- | --- | --- |
| DMN.LP (R) (47,-67,29) – SN.rPFC (R) (32,46,27) | T(39) = -2.93 | 0.005602 | 0.344382 |
| EN.LPFC (R) (41,38,30) – SN.rPFC (R) (32,46,27) | T(39) = -2.74 | 0.009317 | 0.344382 |
| DMN.mPFC (1,55,-3) – SN.rPFC (L) (-32,45,27) | T(39) = -2.67 | 0.010910 | 0.344382 |
| DMN.LP (R) (47,-67,29) – SN.ACC (0,22,35) | T(39) = -2.55 | 0.014713 | 0.344382 |
| SN.rPFC (R) (32,46,27) – SN.SMG (R) (62,-35,32) | T(39) = 2.51 | 0.016399 | 0.344382 |
| DMN.LP (R) (47,-67,29) – SN.rPFC (L) (-32,45,27) | T(39) = -2.41 | 0.020693 | 0.357990 |
| EN.PPC (L) (-46,-58,49) – SN.rPFC (R) (32,46,27) | T(39) = -2.35 | 0.023866 | 0.357990 |
| EN.LPFC (L) (-43,33,28) – SN.rPFC (R) (32,46,27) | T(39) = -2.24 | 0.030912 | 0.405719 |
| DMN.PCC (1,-61,38) – DMN.LP (R) (47,-67,29) | T(39) = 2.03 | 0.049138 | 0.548599 |

**Figure 1.**
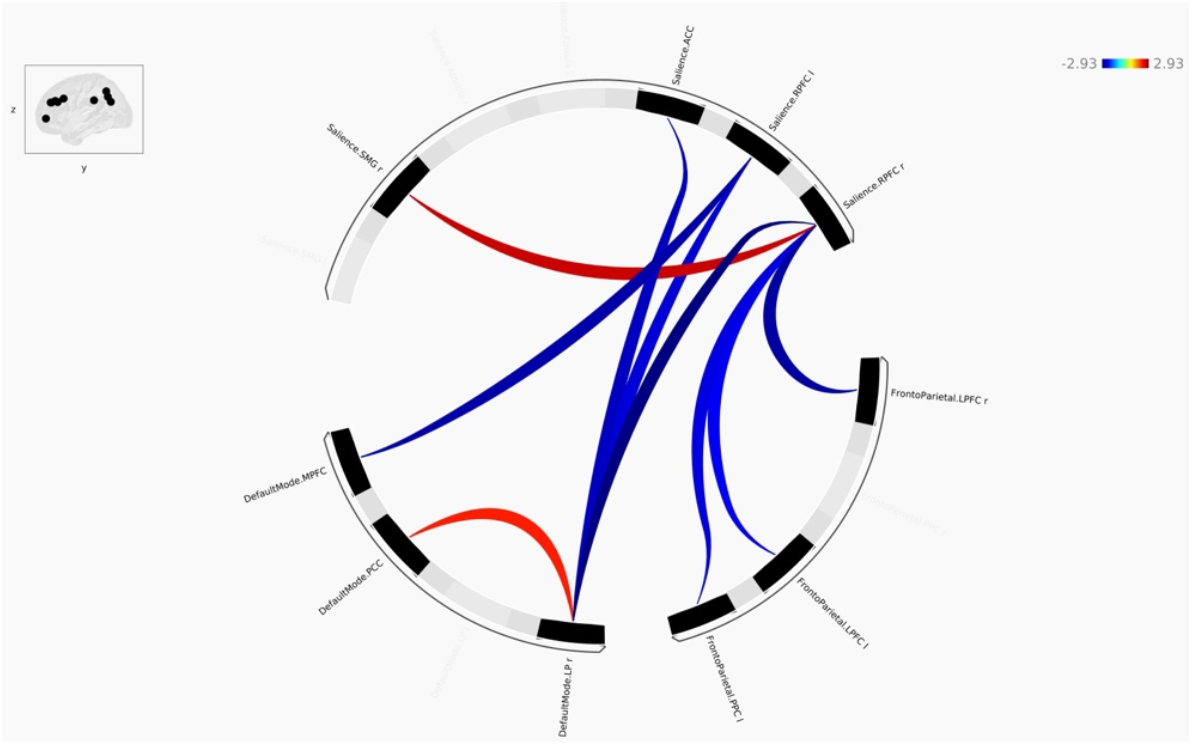
ring picture generated in CONN toolbox. The picture showing uncorrected results of ROI-to-ROI analysis with the network connections that did not pass the FDR threshold (p < 0.05). Red lines indicate increased connectivity, blue lines indicate decreased connectivity. Abbreviations: ACC, Anterior Cingulate Cortex; AI, Anterior Insula; l, left; LP, Lateral Parietal Cortex; LP; Lateral Prefrontal Cortex; MPFC, Medial Prefrontal Cortex; PCC, Posterior Cingulate Cortex; PPC, Posterior Parietal Cortex; r, right; RPFC, Rostral Prefrontal Cortex; SMG, Supramarginal Gyrus.

### Degree centrality

We detected no significant interactions on the voxel level and no significant interactions on the ROI level (see Supplementary Table S1).

## DISCUSSION

In the present study, we tested whether musical creativity training in DHP was associated with changes in seed-based and ROI-to-ROI connectivity pattern or degree centrality within the DMN, EN, and SN. Contrary to our hypothesis, no statistically significant group-by-time interaction effect was observed to show divergent trends between the active and control groups across these networks, despite the previously revealed changes in behavior and task-related brain activities.

The findings are consistent with previous studies reporting limited or non-significant effects of behavioral or cognitive training on intrinsic FC (Cousijn et al., 2014; Tseng et al., 2019). Significant rsFC changes were not observed after verbal creativity training in adolescents, in the seeds including DMN and EN, which were significantly activated during creativity task compared to control task (Cousijn et al., 2014). Working memory training RCT in children showed little evidence of significant differences in intra- or inter-network rsFC between active and control groups across major intrinsic networks including DMN and SN (Tseng et al., 2019). Also, a recent pilot study of clinical trial using rs-fMRI (Wu-Chung et al., 2025) showed that a 6-week music cognitive program in elderly people did not alter overall network indices. Furthermore, a narrative review of exercise and mindfulness interventions reports that many rigorous interventional studies produce minimal or inconsistent rsFC effects, highlighting that null training effects on rsFC are not unusual (Wing et al., 2025). These findings can be interpreted as indicating that large-scale resting-state networks may be relatively stable traits, or that experience-dependent plasticity may preferentially manifest during task engagement rather than during unconstrained rest.

In contrast, as mentioned in the introduction, some fMRI studies have reported training-related changes in rsFC, particularly following verbal creativity training interventions (Wei et al., 2014; Fink et al., 2018; Sun et al., 2020), often involving task-relevant regions or networks. Such discrepancies may reflect differences in training intensity/length, participant age and engagement, network specificity, or analytic sensitivity related to specific network metrics. For example, the interventional methods used in studies by Fink et al. (2018) or Sun et al. (2020) were focused on verbal creativity training conducted over weeks, while musical creativity workshop in our study was held for 2 days.

Taken together, these results suggest that while creativity training can potentially modulate rsFC, such effect may be more consistently detectable during task engagement or under conditions that directly recruit trained processed, rather than during rest. Thus, the present findings refine models of musical creativity training-induced plasticity and guide future research direction by delineating the conditions under which changes in intrinsic network organization emerge.

## FUNDING

This study was supported by OpenAccess grant of Czech-Bioimaging (LM2015062 and LM2018129) to the MAFIL core facility of CEITEC, MUNI, Brno. The Faculty of Education, Palacký University Olomouc, supported student travel grants and scholarships.

## ACKNOWLEDGMENTS

N/A

## Notes

### Competing Interest Statement

The authors have declared no competing interest.

